# *B*_1_^+^-correction of MT saturation maps optimized for 7T *postmortem* MRI of the brain

**DOI:** 10.1101/2022.07.12.498197

**Authors:** I. Lipp, E. Kirilina, L.J. Edwards, K.J. Pine, C. Jäger, T. Gräßle, EBC consortium, N. Weiskopf, G. Helms

**Author notes:** Corresponding author: Name Ilona Lipp, Department Department of Neurophysics, Institute Max Planck Institute for Human Cognitive and Brain Sciences, Address Stephanstr. 1a, 04103 Leipzig, Germany. These authors contributed equally to this work (N. Weiskopf, G. Helms).

## Abstract

**Purpose:** Magnetization transfer saturation (MTsat) is a useful marker to probe tissue macromolecular content and myelination in the brain. The increased 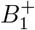 -inhomogeneity at ≥ 7T and significantly larger saturation pulse flip angles which are often used for *postmortem* studies exceed the limits where previous MTsat 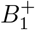 correction methods are applicable. Here, we develop a calibration-based correction model and procedure, and validate and evaluate it in *postmortem* 7T data of whole chimpanzee brains.

**Theory:** The 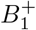 dependence of MTsat was investigated by varying the off-resonance saturation pulse flip angle. For the range of saturation pulse flip angles applied in typical experiments on *postmortem* tissue, the dependence was close to linear. A linear model with a single calibration constant *C* is proposed to correct bias in MTsat by mapping it to the reference value of the saturation pulse flip angle.

**Methods:** *C* was estimated voxel-wise in five *postmortem* chimpanzee brains. “Individual-based global parameters” were obtained by calculating the mean *C* within individual specimen brains and “group-based global parameters” by calculating the means of the individual-based global parameters across the five brains.

**Results:** The linear calibration model described the data well, though *C* was not entirely independent of the underlying tissue and 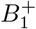. Individual-based and group-based global correction parameters (*C* = 1.2) led to visible, quantifiable reductions of 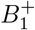-biases in high resolution MTsat maps.

**Conclusion:** The presented model and calibration approach effectively corrects for 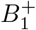 in-homogeneities in *postmortem* 7T data.

## Introduction

Quantitative MRI (qMRI) is a powerful tool to study brain anatomy (1). QMRI parameters in the brain provide measures of tissue myelination, and can be compared between *in vivo* and *postmortem*, across brain regions, across individuals, and even across species, opening the door for a plethora of neuroscience applications (1; 2; 3; 4; 5; 6). In *postmortem* brain, ultra-high field, ultra-high resolution qMRI facilitates studies of white matter myelination and cortical myeloarchitecture across the whole brain with resolutions down to tens of microns (7; 8).

Magnetization transfer (MT) is a contrast mechanism which is particularly specific to brain myelin. The extended lipid–water interface of myelin facilitates efficient magnetization transfer between the MRI-visible water protons (the free water pool) and the MRI-invisible protons of macromolecules (the macromolecular pool) (9; 10). MT imaging exploits the broad absorption line of macromolecular pool protons (1; 9). The broad line allows selective saturation of the macromolecular pool by off-resonance radio-frequency (RF) pulses applied at a frequency remote from the narrow resonance line of the free water (so-called MT pulses). Subsequent transfer of magnetization between the two pools results in a detectable reduction of the measured free water MRI signal (11).

Various acquisition schemes for MT quantification have been developed, which differ in complexity, speed, precision and accuracy (9; 10; 12; 13). One time-efficient approach is multiparameter mapping (MPM) to acquire maps of the MT saturation (MTsat) (13; 14). The MPM approach consists of three RF-spoiled 3D Fast Low Angle SHot (FLASH) acquisitions and provides a high signal-to-noise ratio (SNR) (13; 15; 16); it is therefore particularly suitable for ultra-high resolution qMRI on *postmortem* brains.

While ultra-high field offers advantages with respect to SNR and is therefore a method of choice for ultra-high resolution imaging, it poses particular challenges for quantitative MT mapping. Increased inhomogeneity of the RF transmit field 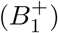 at ultra-high field (17; 18) results in a spatially varying saturation efficiency of the macromolecular pool which is reflected as a bias in the MT estimates which needs to be corrected for (13; 19; 20; 21). The simple bias correction scheme proposed for *in vivo* 3T imaging, where the bias in MTsat is effectively removed by dividing the maps by 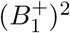 (13), does not work for the high-power MT pulses which are used in ultra-high resolution *postmortem* imaging. *In vivo* work at 3T and 7T demonstrated a correction for 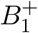-bias on MTsat based on empirical calibration of the MTsat values (20; 22; 23). Here we extend this empirical approach for MT pulses with even higher power for application in *postmortem* MTsat imaging.

To achieve an effective correction, below we empirically determine the dependence of MTsat on 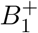 in a calibration experiment on *postmortem* chimpanzee brains. We demonstrate that a simple linear model using only one calibration parameter is sufficient to accurately correct the data. Our correction approach also allows for harmonizing across protocols that apply MT pulses with different flip angles. Thus, it can correct not only for spatial inhomogeneities in 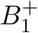 but also account for different imaging protocols or scanner hardware modifications.

## Theory

### Definition of MTsat

In the MT-weighted acquisition in the MPM acquisition scheme a strong off-resonance MT saturation pulse applied every repetition time (TR) causes selective saturation of the macromolecular pool, which is then transferred to the free water pool (Figure 1) (11). The MT saturation (MTsat) is defined as the percentage difference in the free water pool magnetization over one TR period (Figure 1B) relative to the magnetization at the beginning of the TR (13), and is an indirect measure of tissue macromolecular content, *m* (24)^1^.

**Figure 1:**
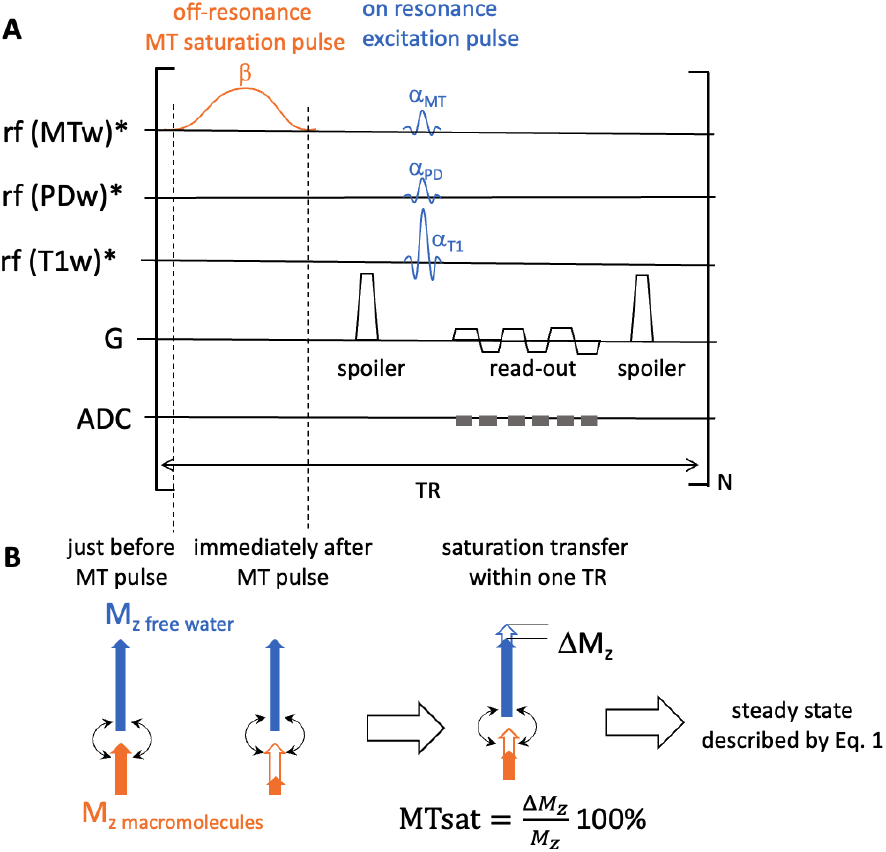
**A** Schematic representation of the MPM pulse sequence for estimation of MTsat, consisting of three 3D FLASH acquisitions, each with a different radiofrequency (RF) excitation scheme (three upper rows). The MT-weighted acquisition uses a low-flip angle on-resonance excitation pulse with flip angle *α*_MT_, and an off-resonance MT pulse with flip angle *β*_nom_ which selectively saturates the macromolecular pool. The PD-weighted and T1-weighted acquisitions use low-flip angle (*α*_PD_) and large-flip angle (*α*_T1_) on-resonance pulses, respectively. Here we assume TR is the same for all acquisitions. **B** MTsat is defined as the difference in the free water magnetization due to the MT pulse over one TR, expressed in percent units of the free water pool magnetization. MTsat results from the exchange of magnetization between the macromolecular pool, which is selectively saturated by the MT pulse, and the free water pool and is dependent on macromolecular tissue content and the degree of saturation of the macromolecular pool.

### Estimation of MTsat

MTsat is estimated using three 3D FLASH acquisitions with different weightings (see Figure 1): (i) an MT-weighted acquisition (*S*_MT_) using a small on-resonance flip angle *α*_MT_ and an off-resonance MT pulse with nominal flip angle *β*_nom_ which selectively saturates the macromolecular pool, (ii) a proton density (PD)-weighted acquisition (*S*_PD_) using a small on-resonance flip angle *α*_PD_, and (iii) a T1-weighted acquisition (*S*_T1_) using a large on-resonance flip angle *α*_T1_. Assuming MTsat, *α*_MT_ and R1 · TR are all small, MTsat can be estimated using (13; 14):

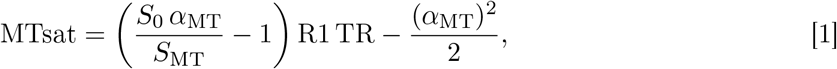

where *S*_0_ is the equilibrium magnetization (proportional to PD) and R1 is the longitudinal relaxation rate. We estimate *S*_0_ and R1 using solutions of an exact algebraization of the Ernst equation^2^ (25):

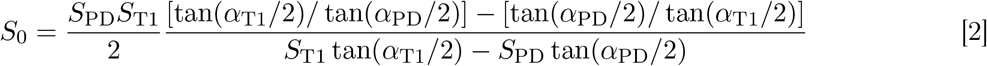

and

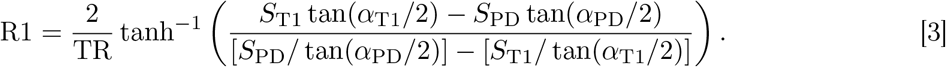

### 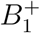 bias in MTsat

Inhomogeneiety of the transmit radio-frequency (RF) magnetic field 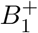 results in spatial variation of the flip angles (*α*_PD_, *α*_T1_, *α*_MT_ and *β*_nom_) across the imaged object such that locally the flip angles are *f*_T_*α*_PD_, *f*_T_*α*_T1_, *f*_T_*α*_MT_ and *f*_T_*β*_nom_, where *f*_T_ is the dimensionless local relative bias in 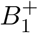 which is determined in a separate calibration experiment (26; 27). For convenience in the following we define

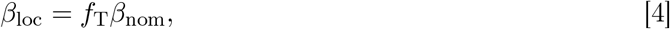

where subscript “loc” denotes the “local” flip angle.

Eqs. [1], [2] and [3] can be be modified to account for the spatial variation of the on-resonance excitation flip angles by substituting *f*_T_*α*_PD_, *f*_T_*α*_T1_ and *f*_T_*α*_MT_ for the respective nominal flip angles. However, Eq. [1] contains an implicit dependence on *β*_loc_ which has not been accounted for: the spatial variation in *β*_loc_ will result in a spatial variation in the saturation of the macromolecular pool, which will result in a spatial variation of the computed MTsat.

MTsat is thus modulated by two factors: the local macromolecular tissue fraction, *m*, and the local MT pulse flip angle, *β*_loc_ (13). While the dependence of MTsat on *m* is an effect of interest used for myelin quantification, its dependence on *β*_loc_ and therefore on 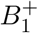 leads to a spatially-dependent bias in MTsat maps which needs to be corrected for to enable accurate whole-brain myelin mapping. Here we present a correction method which aims to render the estimated MT saturation values insensitive to spatial 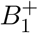 inhomogeneities for *postmortem* brain imaging at 7T, where formaldehyde-fixed tissue and MT saturation pulses with high power are used.

### Dependence of MTsat on the MT flip angle

#### Model assumptions and plausibility arguments

We assume^3^ that the dependence of MTsat on *m* and *β*_loc_ can be simply expressed as a product (28):

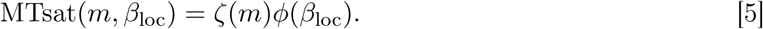

The plausibility of this assumption will be tested experimentally in the Results section. In the following we leave the dependence of MTsat on *m* implicit.

To correct for the effect of 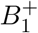 inhomogeneity on MTsat we need to determine the function *ϕ*(*β*_loc_).

Physical considerations reveal that *ϕ*(*β*_loc_) must have a sigmoidal dependence on *β*_loc_ (Figure 2A). At low *β*_loc_, MTsat has a quadratic dependence due to the differential absorption law of the macromolecular pool (9). On the other hand, it approaches a limiting value at very high *β*_loc_ as the macromolecular pool becomes fully saturated after every MT pulse. However, the exact functional form of *ϕ*(*β*_loc_) is in general unknown and is dependent on experimental conditions (magnetic field strength, MT pulse parameters) and characteristics of the investigated tissue (fixation method, temperature).

**Figure 2:**
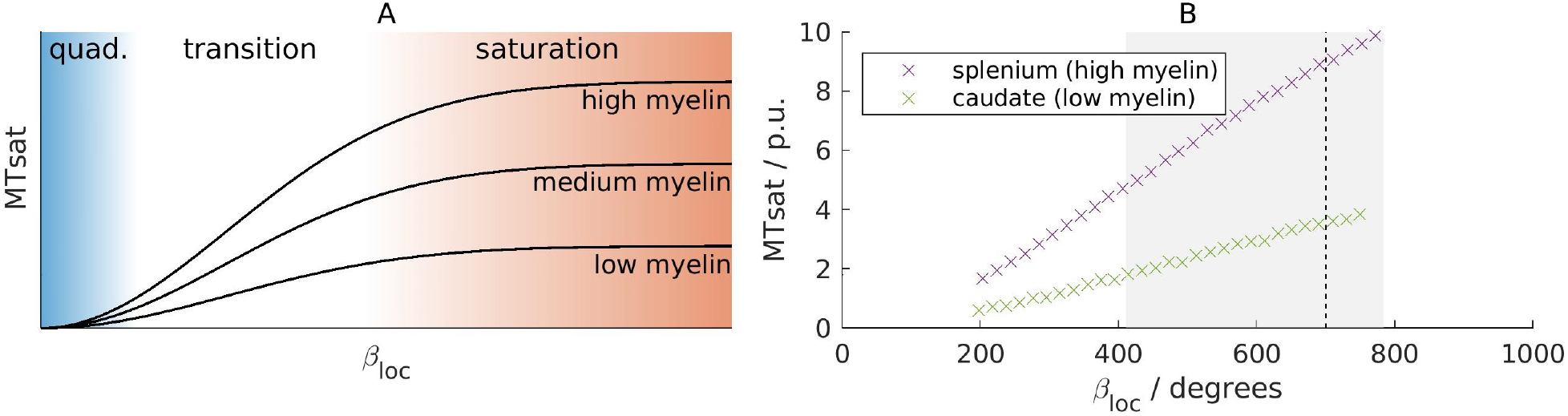
**A:** Schematic representation of the *β*_loc_ dependence of MTsat for different myelination levels (black lines). Three regimes can be distinguished: a quadratic (quad.) scaling regime at low *β*_loc_ (blue), a transition region for intermediate *β*_loc_ (white), and the approach to full saturation of the macromolecular pool at high *β*_loc_ (red). MTsat values corresponding to anatomical regions with different myelination levels are represented by different multiplicative scalings of a universal function of *β*_loc_ (*ϕ*(*β*_loc_); Eq. [5]). **B:** The experimentally observed *β*_loc_ dependence of MTsat in single voxels in two distinct anatomical regions: the highly myelinated splenium of the corpus callosum (purple) and the low myelinated caudate nucleus (green). The gray shaded area shows the range of *β*_loc_ values over the sample when *β*_nom_ = *β*_ref_, the reference MT flip angle (here 700^◦^), which is shown by the dashed line. MTsat shows an approximately linear increase with *β*_loc_ over this shaded area, suggesting that we are in the transition region. Note that the ability to distinguish these two areas increases with increasing *β*_loc_.

To provide a correction of MTsat for a particular experiment we must find a model of the functional form of MTsat(*β*_loc_) for the given experimental conditions over the range of *β*_loc_ observed in the experiment, which we can then use to correct MTsat to a reference MT flip angle, *β*_ref_.

Here we develop an empirical approach to obtain a correction for the special case of large *postmortem* brains (such as humans, apes, whales, elephants) investigated with an ultra-high field MRI scanner. The essence of this approach is to make repeated measurements at a range of *β*_loc_ by scaling *β*_nom_ to obtain a set of MTsat(*β*_loc_) for each voxel (20; 23). These measured data can then be combined to elucidate the *β*_loc_ dependence and derive a correction for it. While we examine the specific case of chimpanzee brains at 7T here, the general empirical approach could be applied to derive a correction model for any case.

### Linear model of the dependence of MTsat on the MT flip angle

Figure 2B shows the empirical dependence of MTsat on *β*_loc_ measured in two distinct regions of a *postmortem* formalin fixed chimpanzee brain. The shaded area gives the range of *β*_loc_ values typically used in *postmortem* experiments. It can be seen that MTsat is empirically linear over the acquired range of *β*_loc_ for both a gray matter (low myelin) and a white matter (high myelin) region. The linear dependence of *β*_loc_ suggests that we are in a linear transition region between the quadratic regime and the saturated regime (Figure 2A). We thus propose a linear functional form for the dependence of MTsat on *β*_loc_:

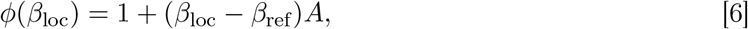

where *A* is a coefficient independent of *m* and *β*_loc_, and *β*_ref_ is the reference MT saturation flip angle we want to map MTsat to. Here we have used the fact that the function *ϕ*(*β*_loc_) is defined up to a multiplicative factor to define *ϕ* such that *ϕ*(*β*_ref_) = 1. Under these assumptions, Eq. [5] can be rewritten as:

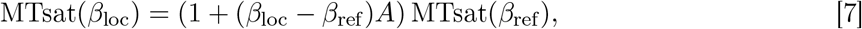

where MTsat(*β*_ref_) is the corrected MT saturation at *β*_ref_. The assumed multiplicative dependence on *ζ*(*m*) (see Eq. [5]) enters this equation through MTsat(*β*_ref_). This model can also be regarded as the first order Taylor series approximation of MTsat(*β*_loc_) around *β*_ref_.

The model parameter *A* can be estimated from the experimental data by voxel-wise linear regression of Eq. [7] using MTsat(*β*_loc_) as the dependent variable and (*β*_loc_ −*β*_ref_) as the independent variable. MTsat(*β*_ref_) is then given by the intercept of the linear model, while the parameter *A* is given by the slope divided by the intercept of the model.

### Correction of MTsat maps using the linear model

Once the value of *A* has been determined in a calibration experiment it can be used to correct the bias in MTsat maps. By rearranging Eq. [7] for MTsat(*β*_ref_) we obtain the transformation of MTsat that corrects it from its value at *β*_loc_ to its value at *β*_ref_ :

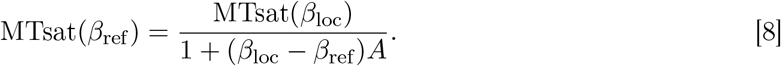

As in Ref. (14), Eq. [8] can be written in terms of a calibration parameter *C* = *β*_ref_ *A*:

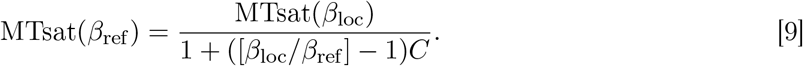

Using Eq. [4] we can write *β*_loc_*/β*_ref_ = *f*_T_*β*_nom_*/β*_ref_ and so

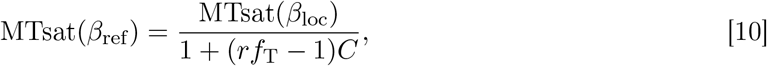

where

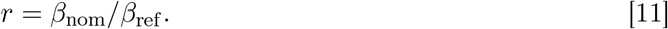

The general Eq. [10] applies for any *β*_nom_ within the range of validity of the model. However, in the usual case where *β*_ref_ = *β*_nom_ (i.e. we have chosen the reference flip angle for the parameter *C* estimation to be the nominal flip angle which we will use for future data sets), *r* = 1 and so Eq. [10] simplifies to

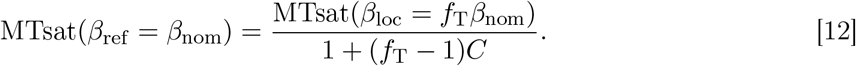

For convenience in the following, we can write Eq. [12] in terms of a local correction factor *F* (*f*_T_, *C*)

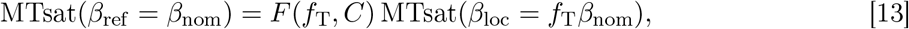

where

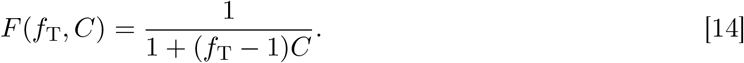

In Eq. [12] the utilized *C* can be the result of fitting to data from each voxel separately, data aggregated across all voxels in a single brain or data aggregated across several brains. Consequently, the correction can be applied with voxel-specific, individual-based global or group-based global parameters to obtain less precise but more robust corrections. Since a generalisation of the correction is desirable, i.e., a globally fixed parameter would be preferred, a comparison between these approaches is made below.

## Methods

### *Postmortem* tissue specimens

Five whole *postmortem* chimpanzee brains (2 females, aged from 12 to 43 years) were studied. These brains were acquired from wild and captive deceased chimpanzees who died unexpectedly of causes not related to this study (see Table S.1 in the Supporting Information for more information). The general preparation of the tissue is described in Refs. (29; 30; 31). *Postmortem* interval before fixation varied from 1 to over 16 hours between brains (Table S.1). Different ages and different *postmortem* intervals before fixation resulted in a broad variation of R1 (between 1.1 and 2.6 s^−1^) and R2^∗^ (between 22 and 37 s^−1^; Table S.1). Thus our sample covers the broad variety of animal ages, tissue fixation conditions and variation of tissue quality typically used in *postmortem* experiments on hominoid brains.

### Data acquisition setup

For MRI data acquisition, the brains were placed in a spherical acrylic container filled with perfluoropolyether (Fomblin). Constant temperature during the scanning (27.5^◦^C to 33.5^◦^C) was facilitated by a warm air stream and monitored by a sensor (LUXTRON Corporation, CA, USA). See Supplementary methods in the Supporting Information for more details.

### MRI acquisition

All data were acquired on a 7T whole-body MRI system (Terra 7T, Siemens Healthineers, Erlangen, Germany) equipped with a 1-channel transmit/32-channel receive RF head coil (Nova Medical, Wilmington, MA, USA).

### Multi-parameter mapping (MPM) protocol

An MPM protocol (14; 15; 32) was used (Figure 1) consisting of ultra-high resolution 3D FLASH images recorded with an isotropic resolution of 300 *µ*m (matrix: 432 *×* 378 *×* 288; readout bandwidth (BW) = 331 Hz/pixel; TR = 70 ms; twelve equidistant echoes (echo times [TE1,…, TE12] = [3.63, …, 41.7] ms); excitation flip angles: *α*_T1_ = 84^◦^, *α*_PD_ = 18^◦^, *α*_MT_ = 18^◦^ for T1-, PD- and MT-weighted images, respectively). A Gaussian-shaped MT saturation pulse at 3 kHz off-resonance with *β*_nom_ = 700^◦^ and length 6 ms gave MT weighting. No partial k-space acceleration was employed.

### Calibration experiment

The calibration experiments were performed using an MPM protocol with similar imaging parameters, but at a lower isotropic resolution of 2.1 mm (matrix: 64 *×* 56 *×* 48, BW = 322 Hz/pixel, [TE1,…, TE12] = [3.6, …, 41] ms) to accelerate the acquisitions.

T1-weighted and PD-weighted images were acquired once at the beginning of each session, followed by MT-weighted images with *β*_nom_ ranging from 220^◦^ to 760^◦^ (in two brains to 700^◦^ due to reaching hardware safety limits), in 20^◦^ intervals. The order of the different acquisitions was pseudo-randomized (see Supplementary methods in the Supporting Information) to balance out potential drifts related to heating or scanner instabilities.

A brain mask was obtained for each brain through intensity thresholding of the first echo of the T1-weighted images.

### 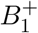 mapping

Maps of the RF transmit field 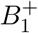 were obtained using the method in Refs. (26; 27) using a spin echo–stimulated echo 3D-EPI sequence (4 mm isotropic resolution, matrix: 48 *×* 64 *×* 48, TR = 500 ms, TE = 40.54 ms; mixing time (TM) = 34.91 ms; spin echo flip angles from 120^◦^–330^◦^ in 15^◦^ increments; GRAPPA acceleration factor = 2 *×* 2) and B0 mapping using a gradient echo sequence (2 mm isotropic resolution, matrix: 96 *×* 96 *×* 64, TR = 1020 ms, TE = 10 and 11.02 ms, excitation flip angle = 20^◦^).

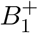 maps were computed with the hMRI toolbox (14) using a global reference T1 = 500 ms. 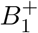 maps were smoothed using Gaussian smoothing (8 mm median filter kernel) and then divided by a brain mask smoothed in the same way. 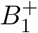 maps were then upsampled to the respective resolution of the FLASH images using FSL `flirt` (using image header information only). *f*_T_ was determined by dividing the obtained relative 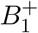 map in percent units by 100%.

### Pre-processing and MTsat calculation

MPM maps were computed separately for each nominal MT pulse flip angle (*β*_nom_).

All weighted FLASH images were corrected for off-resonance related distortions in the readout direction, which alternated between odd and even echoes due to the bipolar readout scheme. First the geometric mean of the first and third echo was calculated, which is an estimate of a virtual second echo image acquired with the opposite readout polarity (33). Using the second acquired echo and the virtual echo as input, the HySCO algorithm of the ACID toolbox (http://diffusiontools.com/) was used to estimate the distortions and correct all acquired echoes (34).

The effective transverse relaxation rate (R2^∗^) was estimated using an ESTATICS (35) weighted log-linear least squares fit, as implemented in the hMRI toolbox (14; 36). No registration was performed between the weighted images. From this fit, the weighted images were extrapolated to TE = 0. These images were then used to calculate MTsat (13) using Eqs. [1], [2] and [3]. In contrast to Ref. (13), we used the local excitation flip angles in the calculation and not the nominal flip angles.

For the analysis with the low resolution calibration data, we created binary masks excluding areas strongly affected by air bubbles. We did this by fitting a simple ordinary least squares model to describe the signal decay over the echos with an exponential function. Voxels in which the model explained less 95% of the variance were excluded from the statistical analysis of the calibration data.

### Calibration parameter estimation and correction

For each voxel within the brain mask and each *β*_nom_ the local *β*_loc_ for the calibration experiment was calculated using Eq. [4] and the experimental *f*_T_ map. The experimental dependence of MTsat on *β*_loc_was fit voxel-wise using Eq. [7] using nonlinear regression as implemented in Matlab’s nlinfit (R2021a). The fit parameters provide a voxel-wise estimation of the calibration parameter *A* in Eq. [7]. nlinfit also gave an estimate of the standard errors of the parameters. The parameter *C* in Eq. [12] was obtained by multiplying by the nominal target flip angle *β*_ref_ (700^◦^ converted to radians). Similarly, the standard error of the fitted parameter *C* was obtained by multiplying the standard error of *A* by the nominal target flip angle *β*_ref_ (700^◦^ converted to radians) and converted to % units relative to the estimated *C*.

When the *β*_loc_ is too low, MTsat cannot be estimated reliably (23). We therefore excluded data points with *β*_loc_ less than our lowest nominal flip angle (220^◦^) from the fit. Low SNR and artefacts can give rise to negative MTsat estimates; these data points were also excluded.

Descriptive statistics of *C* were calculated across all voxels in the brain mask. The mean of each calibration parameter across each specimen was used as an individual-based calibration parameter for this specimen. Then, the means of these individual-based parameters across all specimen were used as group-based calibration parameters (a group-based *C* = 1.2 for *β*_nom_ = 700^◦^ was determined). Individual-based parameters were rounded to two decimal places before applying bias correction. Finally, two different corrections were applied to high-resolution MTsat maps, using the maps and the two different types of calibration parameters (individual-based vs group-based).

### Data and code availability

The analysis code can be found at https://github.com/IlonaLipp/MTcalibration. Data associated with this manuscript can be found at https://doi.org/10.17617/3.803OMM.

## Results

### The calibration coefficient *C*: estimation uncertainty, within- and between-brain variation

Examples of the MTsat dependence on *β*_loc_ obtained in the calibration experiment for individual voxels are shown in Figure 3. Plotted in normalized coordinates the dependencies measured for different voxels and different brains overlapped within experimental error, supporting the plausibility of our model assumption of a universal dependence of MTsat on *β*_loc_. The dependence of MTsat on *β*_loc_ was very well described by the proposed linear model for all brains (Table 1, Figure 3, Figure 4) as reflected in the high goodness of fit, with the average (median) *R*^2^ across the brain exceeding 0.95 in all five specimens (Table 1). The uncertainties of *C* estimated for each voxel and averaged across the brain lay between 0.6–1.1% of the mean value with the exception of brain 1 with an average uncertainty of 3.2% due to the scanner drift during the calibration experiment for this brain (Table 1, Figure 4). Larger errors in the estimation of *C* were observed at the edges of the brain (indicated by the large SD of the model fit in Figure 4) and around air bubbles within the sample, probably due to the low signal and the effect of mechanical and scanner drifts during the calibration experiment.

**Table 1:** Estimated values for the calibration parameter *C* and related variances. The mean, between voxel variance and the within-voxel uncertainty of the estimated parameters *C* across all voxels for the five brains are provided. Voxels with values of *C <* 0 (assumption that only positive correlations are physical) or *C >* 1.4 (assumption that MTsat(*β*_loc_) is positive for all *β*_loc_ down to 220^◦^ given *β*_ref_ = 700^◦^; derived from Eq. [10]) were excluded from these statistics. Uncertainties and variance are provided in % from *C*. The within-voxel uncertainty was estimated from the SD of the fitted parameter and is reported in % from the fitted *C*. The between-voxel variance was estimated as the SD across the brain converted to % of mean *C*. The quality of the linear model fit (coefficient of determination *R*^2^) is provided as mean *±* SD across the brain. To quantify the dependence of the estimated *C* on the underlying 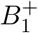 and the corrected tissue MTsat, whole-brain voxel-wise partial Spearman correlation coefficients were calculated.

| Brain | $C$ | between<br>voxel<br>variance of<br>$C$ (% from<br>mean $C$) | within-voxel<br>uncertainty of<br>$C$ (% from<br>fitted $C$) | model fit | $\rho_{\text{part}}(\text{MTsat})$<br>$C$ vs $B_1^+$ | $\rho_{\text{part}}(B_1^+)$<br>$C$ vs<br>MTsat |
| --- | --- | --- | --- | --- | --- | --- |
| brain 1 | 1.242±0.071 | 5.733 | 3.233±23.160 | 0.952±0.096 | −0.124 | −0.635 |
| brain 2 | 1.177±0.030 | 2.567 | 0.643± 0.526 | 0.996±0.013 | −0.503 | −0.534 |
| brain 3 | 1.207±0.046 | 3.802 | 0.743± 0.978 | 0.993±0.029 | −0.390 | −0.622 |
| brain 4 | 1.191±0.047 | 3.951 | 1.149±10.634 | 0.988±0.042 | −0.369 | −0.574 |
| brain 5 | 1.227±0.053 | 4.332 | 0.960± 1.841 | 0.990±0.039 | −0.312 | −0.747 |

**Figure 3:**
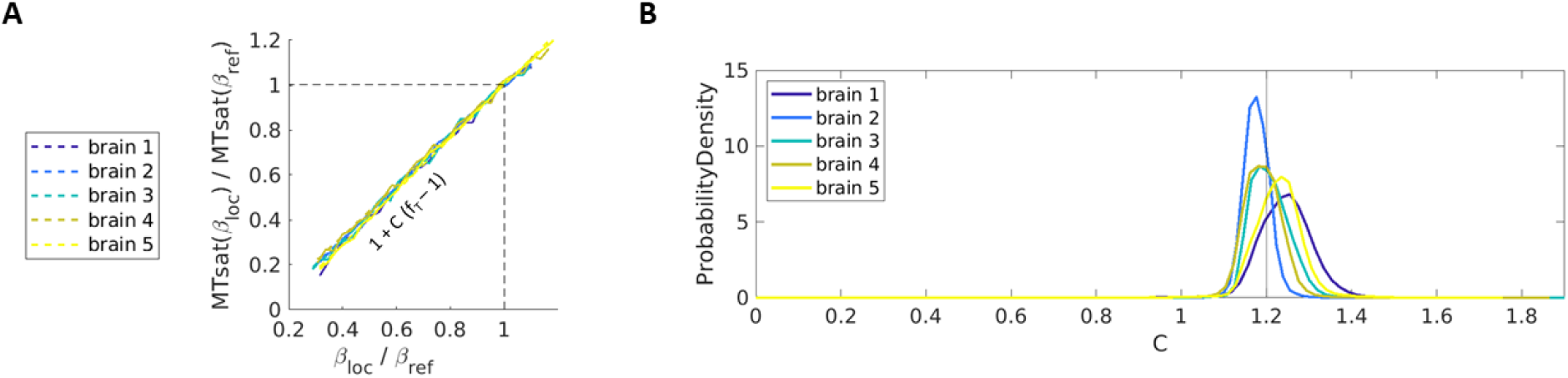
Distribution of the calibration parameter *C*. **A**. Voxel-wise dependencies of MTsat on *β*_loc_ obtained in the calibration experiment for exemplary white matter voxels from five brains (solid lines) together with linear model fit (dashed lines). The dependencies are presented in the normalized coordinates MTsat(*β*_loc_)*/*MTsat(*β*_ref_) and *β*_loc_*/β*_ref_. The linear model yielded high goodness of fit, providing voxel-wise estimation of *C*. In the normalized coordinates *C* corresponds to the slope of the fitted linear dependence. Note that experimental data from all brains nearly overlapped when plotted in normalized coordinates and are therefore described well by similar values of *C*. **B**. Histograms for *C* across all voxels of each specimen. The histograms obtained for five brains overlapped and were centered around *C* = 1.2 (gray line), with some variation within and between the brains.

**Figure 4:**
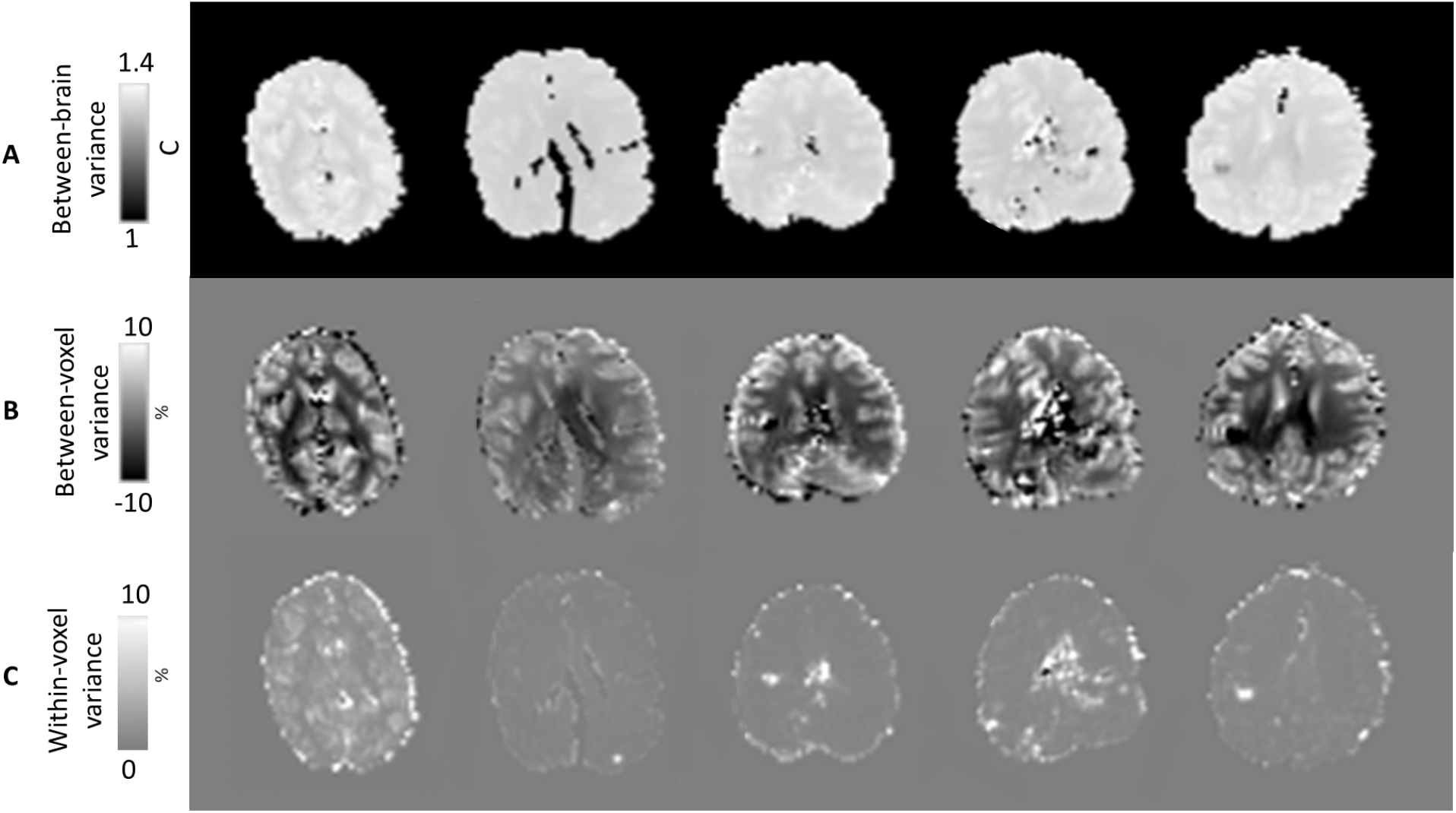
Variance of the calibration coefficient *C*. **A** Maps of calibration coefficient *C* for the five investigated brains obtained by a voxel-wise fit of the experimental dependence of MTsat on *β*_loc_. All five brains demonstrated similar values of the calibration coefficient *C* all close to 1.2, with very low variation across the brain. **B** Between-voxel variance in *C* is illustrated through % difference from the mean value. **C** Maps of within-voxel uncertainty for *C* estimates, expressed in the parameter SD converted to % to *C* of that voxel.

The means of the calibration coefficient *C* for the five brains were all close to 1.2 (varying between 1.18 and 1.24), with a mean of 1.209 and an SD of 0.026 between the brains corresponding to 2% of the mean *C* value. The histograms of *C* for the five brains showed unimodal overlapping distributions (Figure 3). The between-voxel variation (standard deviation; SD) of *C* within each brain ranged from 2.6–5.7% of the mean value and exceeded the averaged within-voxel uncertainty of the *C* and between brain variation of *C* (Table 1).

That we obtained similar values of the calibration coefficient for brains with different ages and varying fixation conditions demonstrates the wide generalizability of the proposed calibration approach for *postmortem* brain imaging.

### Residual tissue type dependence of *C*

A key assumption behind the applied approach was that *C* is independent of the macromolecular content and tissue type (Eq. [5]). This assumption was largely supported by the experimentally obtained values of *C* (Figure 3), which showed very close agreement across the brains and even between the brains with different fixation conditions and from individuals of different ages. However, the spatial distribution of *C* demonstrated residual systematic contrast between gray and white matter regions, with white matter showing on average 2.5% lower values of *C* as compared to gray matter (Figure 4). To illustrate the residual dependence of *C* on tissue type and underlying 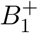, we computed the histograms of *C* within cortical and white matter voxels separately in one of the brains (Figure 5). These show different distributions of 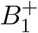 and *C* between cortex and white matter. Also, *C* showed small systematic differences between areas of low and high 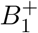. Figure 5B shows that the effects of 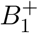 and tissue type are partly independent of each other, as the relationship between 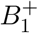 and *C* is clearly visible even if just white matter voxels are considered. Additional analysis of this residual dependence using Spearman correlation is presented in Supplementary residual *C* dependence on tissue type and 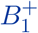 in the Supporting Information.

**Figure 5:**
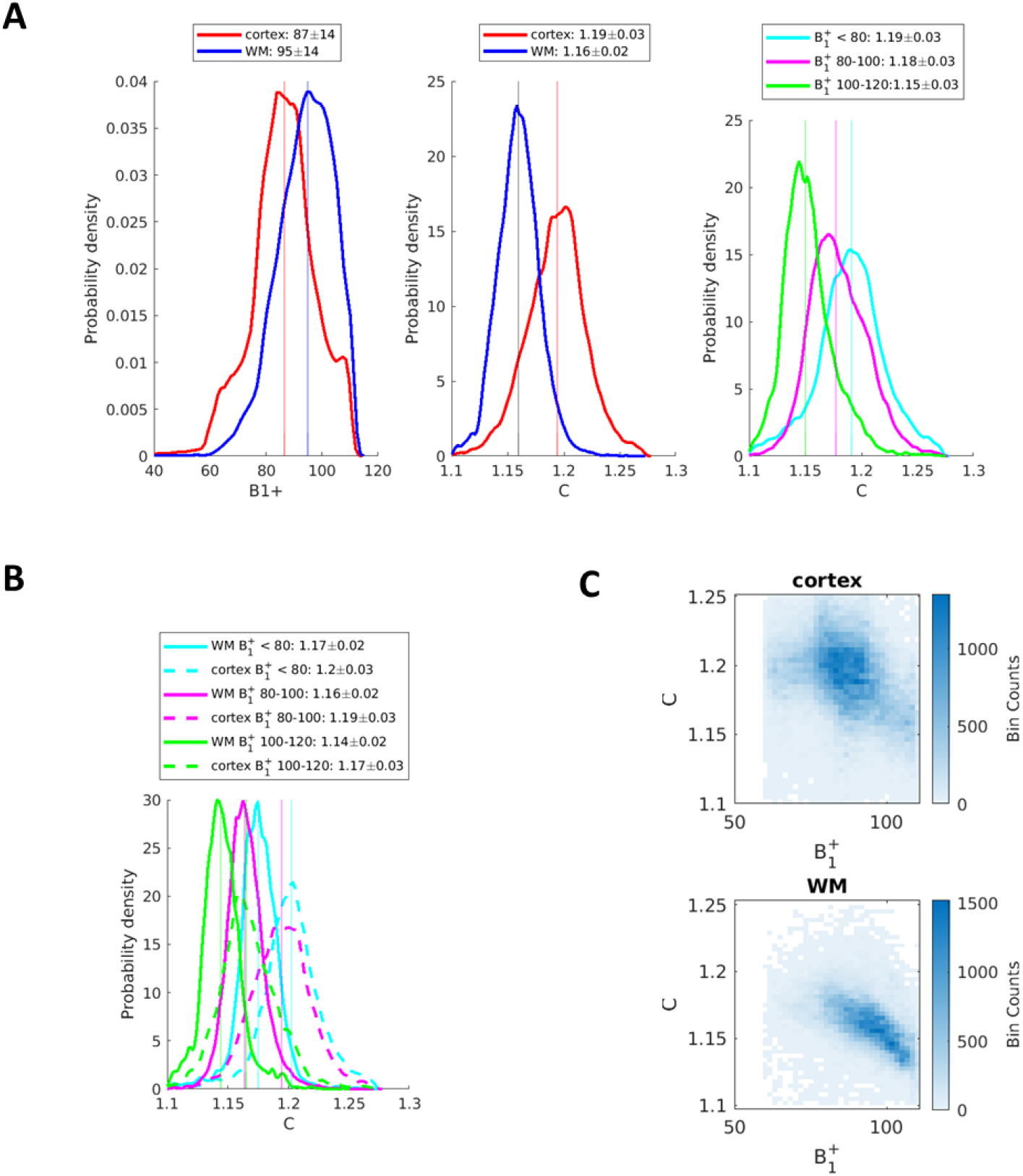
Effects of tissue type and 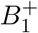 on *C* in a representative brain (specimen 2). **A** Left: Histograms show lower values of 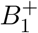 in cortex than in white matter (WM). Middle: Parameter *C* is higher in cortical voxels than in WM voxels. Right: Distributions of *C* based on 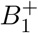, showing the lowest values of *C* in regions with the highest 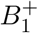. Both tissue type and 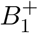 independently explain variance in *C* across the brain, as quantified by partial correlation coefficients (Table 1). Median *±* interquartile range are provided in the legend. **B** shows the interaction between tissue type and 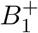. The effect of 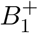 (denoted by different colors) can be seen for both tissue types (WM in solid line, cortex in dashed line). **C** The relationship between 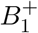 and *C* is additionally illustrated in density scatter plots for cortex and WM voxels separately. The relationship is particularly visible in the WM. Cortex and white matter masks were obtained using Freesurfer (https://surfer.nmr.mgh.harvard.edu/).

Next, we tested the practical relevance of this residual dependencies by comparing the bias correction with the voxel-wise vs average *C* values.

### Bias correction in MTsat maps: comparing calibration approaches

We tested the performance of the proposed calibration approach for the correction of the bias in MTsat maps resulting from the inhomogeneity of 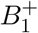. For each brain we corrected the bias in MTsat maps in three different ways, “voxel-wise”, “individual-based” and “group-based”. Voxel-wise refers to using each voxel’s estimated *C* parameter, and individual-based refers to using the specific brain’s median calibration parameter *C* (as reported in Table 1). Group-based refers to correcting with a fixed set of calibration parameters, i.e, the mean across all brains (*C* = 1.2).

Given the residual spatial dependence of *C*, in theory, corrections with voxel-wise *C* should provide most accurate results, while requiring the time-consuming calibration experiments which are only feasible for low-resolution MTsat maps, since repeating the whole calibration experiments for lengthy high resolution scans would require infeasibly long scanning times. Corrections with whole-brain values also requires calibration experiment on each sample, but would be feasible for ultra-high resolution data by performing a calibration experiment at low resolution. Correction of group-based mean values is less accurate but most time efficient, since it requires a calibration experiment for only a subset of representative samples. In the following we evaluated the difference between these three proposed approaches.

### Voxel-based vs. individual-based correction

Figure 6 shows the comparison between voxel-wise and individual-based correction in one exemplary brain. Visible reduction of the bias was achieved with both approaches, with the average difference between corrected maps lying within the *±*0.5 p.u. interval.

**Figure 6:**
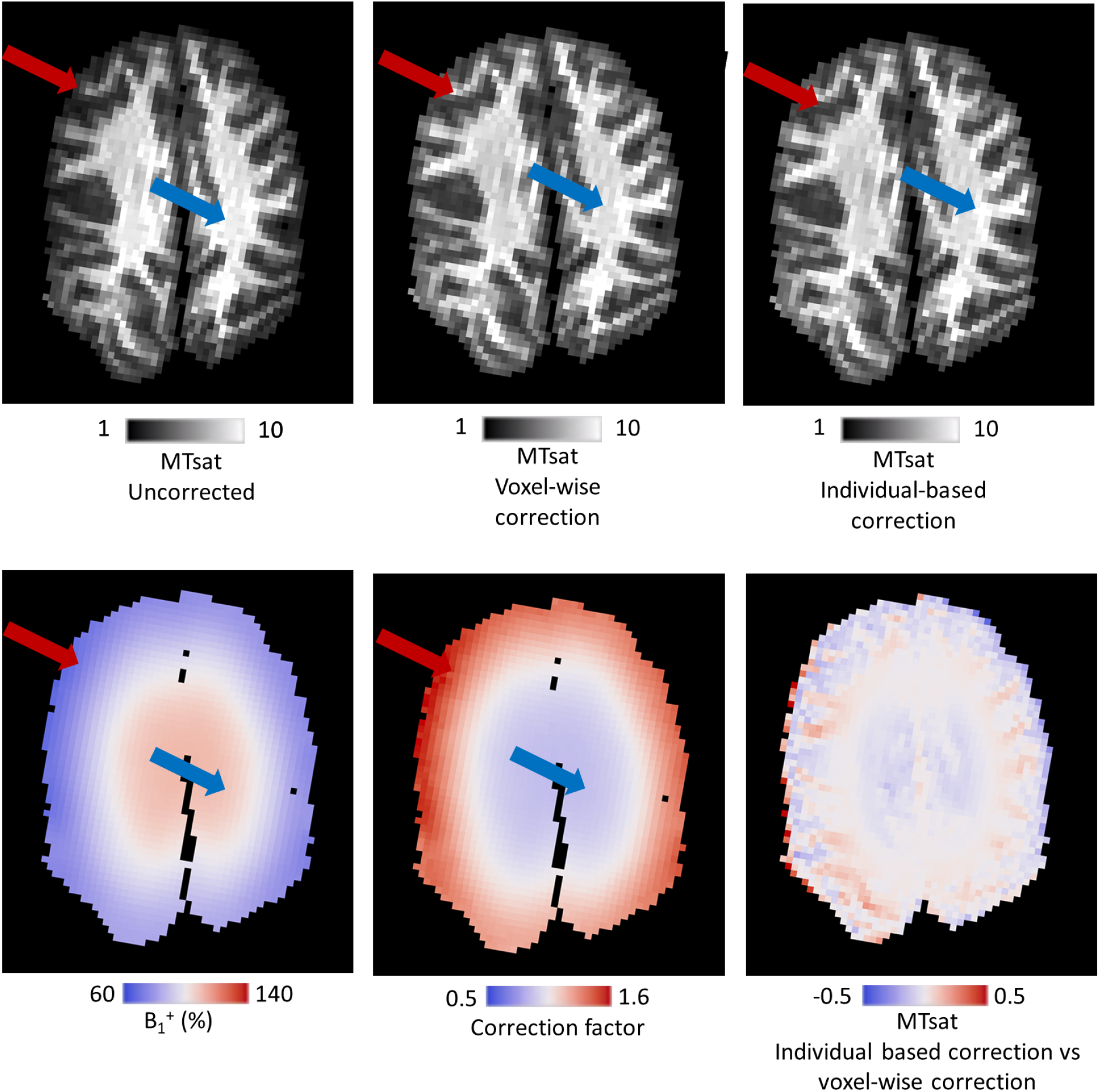
Example maps of MTsat. **Top:** Correction of the MTsat map. An axial slice of the uncorrected map is shown and compared to a voxel-wise correction and the individual-based correction approach. The red arrow indicates a region of low *β*_loc_, whose hypointensity was corrected. The blue arrow indicates an area of high *β*_loc_, whose hyperintensity was corrected. **Bottom:** The *β*_loc_ and corresponding correction factor (calculated using Eq. [14], with either voxel-wise or individual-based mean *C*) are shown, along with the difference map between voxel-wise and individual-based correction. Tissue contrast and features of *β*_loc_ are visible in this map.

To quantify the 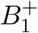 bias and the effect of the correction, spatial Spearman correlations were calculated between the apparent MTsat and 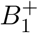 across the brain. Due to field focussing in head-sized objects in ultra high field MRI 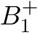 has a maximum at the center of the brain (17; 18). Therefore the 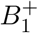 distribution is spatially correlated with anatomy (white matter in the middle of the brain, gray matter in the periphery). We regressed out the effect of anatomy by using the 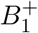-independent measure R2^∗^ (Table 2). The correlation coefficients for uncorrected data ranged from *r* = 0.422 to *r* = 0.633. All three correction approaches were able to reduce that bias (reducing the correlations to between *r* = −0.116 and *r* = 0.307). The correction had an average effect on MTsat of 11–16% of the uncorrected value, while the difference between correction approaches lay in the range of less than 0.5% (for all numbers see Table 2). Therefore, the observed variability in *C* across the brain is negligible compared to the 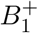-bias in MTsat maps. Additionally, comparing the voxel-wise to the individual-based correction revealed some voxels with very large differences (indicated by the max values reported in Table 2) indicating that voxel-wise corrections fail in some regions prone to artifacts induced by either low signals or scanner instability during the calibration experiment, which can particularly affect brain edges.

**Table 2:**
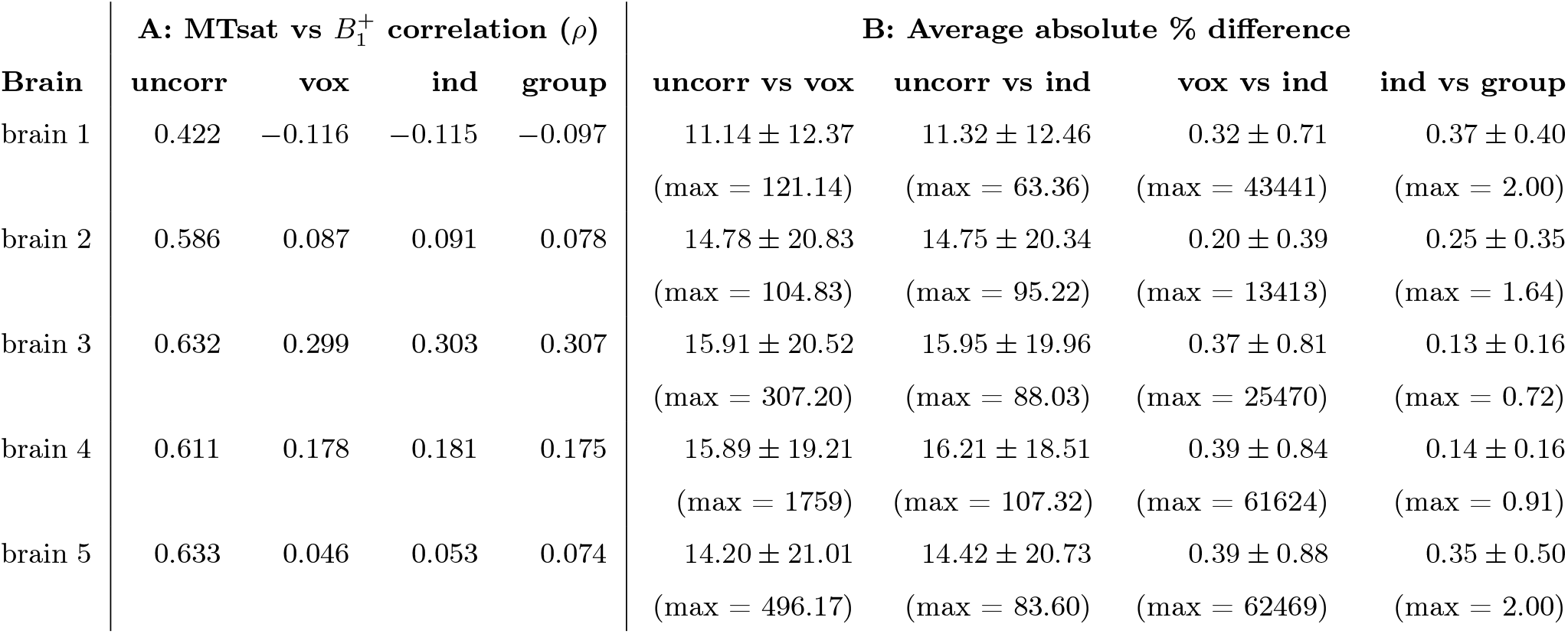
Evaluation of the bias correction in the low resolution calibration data. **A**: Quantification of the bias in the maps. For each brain, the correlation coefficient between 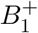 and MTsat across all voxels in the brain was quantified using a partial Spearman correlation coefficient (regressing out R2^∗^), using the uncorrected maps “uncorr”, the maps using the voxel-wise estimated calibration coefficients “vox”, the maps corrected with the individual-based calibration coefficients “ind”, and the maps corrected with the group-based calibration coefficients “group”. **B**: Quantification of the absolute effect of the bias correction. Median *±* interquartile range of the percent change in MTsat due to the correction were calculated across all voxels in the brain and reported in absolute values. We also report maximum absolute difference for the maps. Changes are reported between uncorrected maps and correction with voxel-wise parameters, uncorrected maps and correction with individual-based parameters, between correction with voxel-wise parameters and correction with individual-based parameters, and between the the two global correction approaches.

### Individual-based vs. group-based correction

For each brain we also 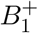-bias corrected the ultra-high resolution MTsat maps in two different ways, “individual-based” and “group-based”, as described above.

In the 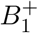-uncorrected data, partial correlation coefficients ranging from *ρ* = 0.440 to *ρ* = 0.589 were observed, reflecting the 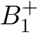-induced bias in MTsat. The relationship between 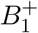 and Mtsat was lower for all corrected data sets and ranged from *ρ* = −0.094 to *ρ* = 0.196 with the individual-based correction and from *ρ* = −0.077 to *ρ* = 0.201 with the group-based correction. We also quantified the effect of the applied corrections on the MTsat values. Across all voxels, the average effect of applying the correction was an absolute change between 11.75 and 18.90% in MTsat (Table 3). Comparing the two correction approaches to each other, the individually- and group-based corrected MTsat maps differed on average by 0.14–0.42%.

**Table 3:** Evaluation of the high-resolution bias correction. **A: Quantification of the bias in the maps**. For each brain, the correlation coefficient between 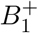 and MTsat across all voxels in the brain was quantified using a partial Spearman correlation coefficient (regressing out R2^∗^), using the uncorrected maps “uncorr” (left column), the maps corrected with the individual-based calibration coefficients “ind” (middle column) and the maps corrected with the group-based calibration coefficients “group” (right column). **B: Quantification of the absolute effect of the bias correction**: Median *±* interquartile range % in MTsat due to the correction were calculated across all voxels in the brain. Absolute changes are reported between uncorrected maps and correction with individual-based parameters (left column), between uncorrected and correction with group-based parameters (middle column), and between the the two correction approaches (right column).

| Brain | A: MTsat vs $B_1^+$ correlation ($\rho$) | | | B: Average absolute % difference | | |
| --- | --- | --- | --- | --- | --- | --- |
|  | uncorr | ind | group | uncorr vs<br>ind | uncorr vs<br>group | ind vs<br>group |
| brain 1 | 0.440 | -0.087 | -0.069 | 12.17 ± 9.33 | 11.75 ± 8.95 | 0.39 ± 0.38 |
| brain 2 | 0.523 | 0.018 | 0.004 | 18.33 ± 15.95 | 18.74 ± 16.39 | 0.31 ± 0.34 |
| brain 3 | 0.556 | 0.196 | 0.201 | 17.28 ± 13.67 | 17.10 ± 13.50 | 0.14 ± 0.15 |
| brain 4 | 0.589 | 0.099 | 0.092 | 18.70 ± 16.52 | 18.90 ± 16.75 | 0.16 ± 0.14 |
| brain 5 | 0.469 | -0.094 | -0.077 | 17.19 ± 14.28 | 16.67 ± 13.76 | 0.42 ± 0.43 |

Figures 7A and B illustrate that bias correction of the MTsat values across the brain visibly reduces the bias and yields more distinct histogram peaks of gray and white matter. Figure 7C shows that uncorrected cortical surface MTsat maps strongly resemble the 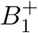 distribution. After correction the maps show patterns reflecting myeloarchitecture, with the highly myelinated primary cortical areas standing out. For example, the expected high myelination of primary motor and primary somatosensory cortex along the central gyrus only becomes apparent after the correction. This demonstrates that bias correction in MTsat maps is a crucial step when studying cortical myelination across the entire brain and enables quantitative comparison of myelination between cortical areas.

**Figure 7:**
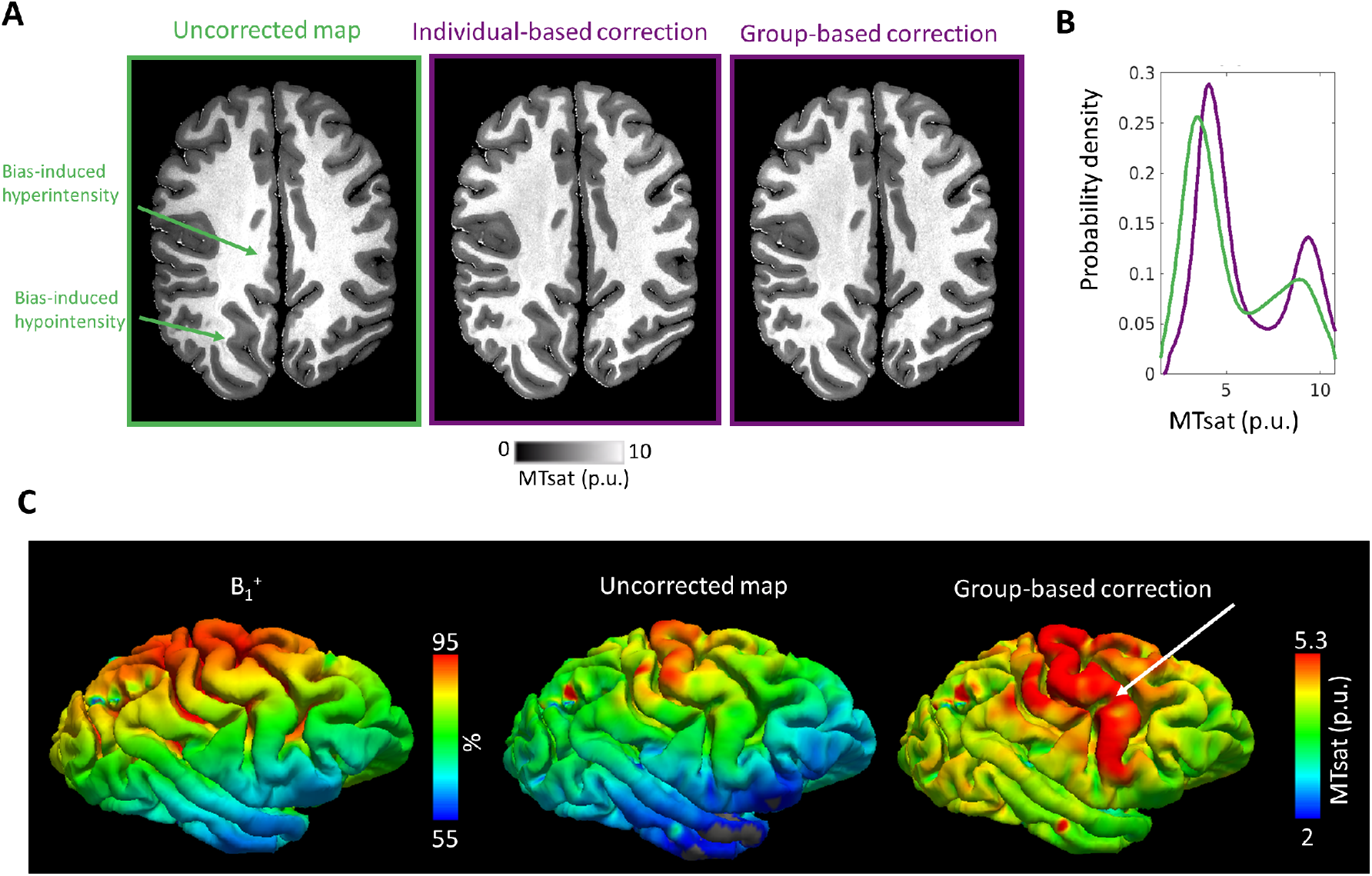
Correction of high resolution images. **A**. The effect of 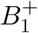 bias correction on an example high resolution MTsat axial slice from brain 2. **B**. A histogram comparing the distribution of the uncorrected map (green) to correction with the individual-based parameters and group-based parameters (purple). The corrected maps show an emphasized bimodal distribution of values, reflecting gray and white matter, whilst the uncorrected map provides a poorer distinction. The bias visible in the uncorrected map is reduced in both corrected maps. The probability density plot was created with the Matlab function `ksdensity` using 100 bins and excluding outliers (data points below the 2nd or above the 98th percentile). **C**. The distribution of 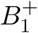 (left) on the Freesurfer (https://surfer.nmr.mgh.harvard.edu/) mid-cortical surface of brain 2 is shown, demonstrating that 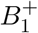 can also systematically vary across cortical regions. If not corrected, this is also reflected in the cortical MTsat maps (middle). Our correction eliminates this bias, revealing the highly myelinated primary cortical areas (right; arrow).

## Discussion

We have developed a calibration correcting for 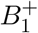-induced biases in MTsat maps and demonstrated its efficacy. It extends previous calibration approaches to the higher 7T static magnetic field strength and the stronger MT saturation pulses used for *postmortem* imaging. A high goodness of fit of the calibration model to experimental data supports the theory-based correction approach. Although the status and quality of the tissue varied significantly, we found a single fixed calibration parameter that significantly reduced the 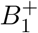-induced bias in all *postmortem* data sets, corroborating the generalizability. This simplifies the implementation as a standard tool, since calibration parameters do not need to be estimated for each *postmortem* specimen individually, which would require the acquisition of additional reference data for each specimen.

### Factors influencing the estimation of *C*

The presented calibration requires the experimental estimation of the calibration parameter *C*. A strong overall goodness of fit suggests the validity of the theoretical model. The *R*^2^ of *>* 0.97 we obtained for all brains vastly exceeds the average *R*^2^ of 0.20 that was reported in a similar experiment conducted on humans *in vivo*, in which the model was fit over large regions which were assumed to be homogeneous (23). Here, we fit the model and estimate *C* voxel-by-voxel, which, unlike region-based analyses, captures spatial inhomogeneities of the calibration curve. However, voxel-wise estimations are more sensitive to variations in statistical noise, 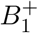 errors and the underlying tissue. This limitation is lifted by using whole brain average or group averaged calibration coefficients.

The calibration approach was based on an assumption that *C* does not depend on 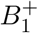, tissue characteristics or tissue type. In our data, the calibration parameters systematically differed between gray and white matter by about 4% (also apparent in a significant spatial correlation between *C* with the corrected MTsat values). Also some residual dependence on 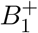 was observed. We calculated global, individual-based calibration parameters for each brain by taking the mean values across all voxels. Corrections using voxel-wise values of *C* did not result in a better correction than using brain-averaged values of *C* (Table 2), so this residual tissue and 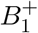 dependence has a negligible effect on correction performance.

### Consistency of *C* across brains

Overall, the variability of the estimated calibration parameters within individual brains (Table 1) was larger than variability between brains. This indicates that factors such as tissue quality, *postmortem* time etc. play only a minor role and a generalized correction is possible. Correction using the group-based mean calibration parameters of our sample (*C* = 1.2) visibly reduced the bias in all brains. Individually optimising the calibration coefficients may yield more accurate correction results. However, when comparing the individually and group-based corrected maps, average differences of less than 0.5% were obtained. In comparison to the average 12–19% change that either correction had on the uncorrected maps, this appears minor. Therefore, a correction with the group-based parameters is recommended when individual-based parameters are not available, i.e., when additional reference data are not available.

For data obtained *postmortem* at 7T with the acquisition parameters used in our study, the following equation with group-based *C* = 1.2 can be applied for correction:

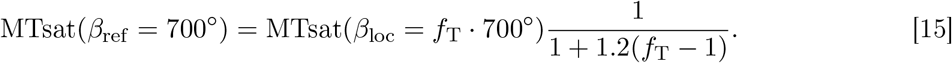

### Correction approach reveals true biological variability

As a ground-truth for MTsat is generally not available, we assessed the performance of the 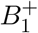-bias correction by directly comparing the different types of corrections with respect to their anatomical validity. All correction approaches had a significant impact on MTsat values, with an average change of up to 20% of the uncorrected value. The two separate gray and white matter peaks in whole sample MTsat histograms became better separated after correction (Figure 7). This indicates that non-biological sources of variance were being reduced. Additionally, cortical surface projections of uncorrected MTsat maps lead to visible biases that correspond to the 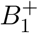 distribution across the cortex (Figure 7). Visual assessment of corrected and original uncorrected MTsat maps showed reduced bias and clearer delineation of anatomical structures (e.g. motor and somatosensory cortex; Figure 7).

### Previous simulation based approach

We used an empirical approach to determine a functional form for the correction of 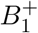 bias. The functional form could alternatively be elucidated by using forward models of the MT effect to simulate the dependence of MTsat on *β*_loc_ and *m* (21). However this method is limited by the need for reliable forward model parameter estimates (exchange times, pool size ranges and relaxation times/lineshapes of the macromolecular and free pools). These are sensitive to tissue preparation methods, difficult to measure, and not generally available.

### Limitation: potential acquisition protocol dependence

Although a linear dependence on the flip angle is expected in the transition region, this specific calibration approach was only tested on one hardware setup and protocol. We show in the Supporting Information how Eq. [15] can be modified (see Eqs. [S.1] and [S.2]) to map the measured MTsat to different *β*_ref_ flip angles within the region where the linear model applies.

However, if data are obtained with major changes in the acquisition protocol (e.g. *β*_nom_ far outside the experimental range used in the calibration), then recalibration may be required using the described calibration experiment. The calibration experiment and estimation of *C* can be performed on a small number of *postmortem* specimens, and the resulting group-based calibration parameters can then be used to correct all additional specimens. Future research may investigate which sequence parameters have no or little impact on the calibration parameters.

## Conclusion

We developed a 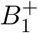 correction of MT saturation MTsat maps using a calibration approach. It extends previous calibration approaches and enables quantitative *postmortem* MT mapping using high power MT saturation pulses at 7T. We showed that a single correction coefficient can visibly reduce 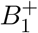-related biases in low resolution data, high resolution data and on cortical myelination maps.

## Acknowledgements

This project has been funded by the Max Planck Society and by the European Research Council under the European Union’s Seventh Framework Programme (FP7/2007-2013) / ERC grant agreement n^◦^ 616905. NW received funding from the European Union’s Horizon 2020 research and innovation programme under the grant agreement No 681094. This project received funding from the Deutsche Forschungsgemeinschaft (DFG, German Research Foundation) – project no. 347592254 (WE 5046/4-2 and KI 1337/2-2). GH was funded in part by the Swedish Research Council, grant NT 2014-6193. IL has been funded by the Deutsche Forschungsgemeinschaft (DFG, German Research Foundation) – project no. 446291874.

We are grateful to Caroline Jantzen, Felix Büttner, Niklas Alsleben and Franziska Zahn for their help with sample preparation and scanning. IL is grateful to Dr Daniele Agostini for being a rubber duck.

## Supporting Information

### Tissue specimens

**Table S.1:** Tissue specimens. Five whole brains from chimpanzees were studied. Sex, age at death, *postmortem* time, cause of death, brain averaged (median *±* IQR) R1 and R2^∗^ values are reported.

| Specimen | Sex | Age | *Postmortem* time | Cause of death | R1 / s<sup>-1</sup> | R2* / s<sup>-1</sup> |
| --- | --- | --- | --- | --- | --- | --- |
| 1 | F | 13 years | 15 hours | poaching | 2.56 ± 0.93 | 22.22 ± 17.69 |
| 2 | F | 43 years | 1 hour | during surgery | 1.08 ± 0.37 | 37.34 ± 24.83 |
| 3 | M | 16 years | < 16 hours | bacterial septicemia | 1.90 ± 0.94 | 35.28 ± 17.71 |
| 4 | M | 15 years | < 12 hours | bacterial septicemia | 2.11 ± 0.70 | 33.26 ± 18.21 |
| 5 | M | 12 years | 6 hours | conspecific aggression | 1.78 ± 0.49 | 24.85 ± 15.59 |

### Supplementary methods

#### *Postmortem* acquisition setup

For MRI data acquisition, the brains were placed in a spherical acrylic container filled with perfluoropolyether (Fomblin) and stabilised in place with sponges. Brains were stored at room temperature for at least two hours before scanning. A warm air stream was directed at the specimen during scanning to achieve constant temperatures closer to normal body temperature. The sample temperature was monitored by a sensor (LUXTRON Corporation, CA, USA) attached to the container surface and ranged from 27.5^◦^C to 33.5^◦^C across the specimens.

#### Order of MT pulse flip angles

The pseudo-random order of MT pulse flip angles during the calibration was: 620^◦^, 300^◦^, 240^◦^, 500^◦^, 400^◦^, 320^◦^, 740^◦^, 520^◦^, 460^◦^, 340^◦^, 280^◦^, 760^◦^, 600^◦^, 680^◦^, 720^◦^, 700^◦^, 560^◦^, 480^◦^, 200^◦^, 640^◦^, 220^◦^, 260^◦^, 540^◦^, 660^◦^, 440^◦^, 360^◦^, 580^◦^, 380^◦^, 420^◦^.

#### Accounting for different reference flip angles without recalibration

Once we have estimated the calibration constant *C* for a given *β*_ref_, MTsat may be scaled to any target flip angle 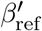 that lies within the range of validity of the model using

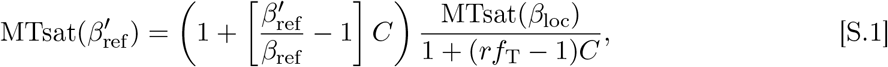

i.e. changing the target flip angle from *β*_ref_ to 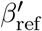 amounts to multiplication of the correction in Eq. [10] by a scaling factor 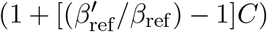. This relation was derived by setting 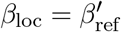 in Eq. [9], solving for 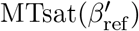, and then inserting MTsat(*β*_ref_) from Eq. [10] into the resulting equation.

In the special case where 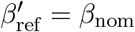 then 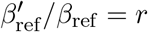 which allows Eq. [S.1] to be written in the same form as Eq. [12]:

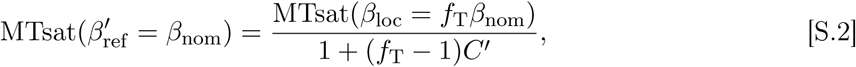

where

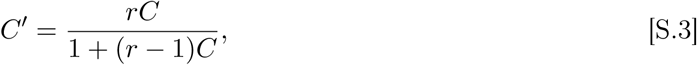

i.e. in this case we can just update the calibration constant to compute the correction to 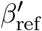 rather than multiplying by a prefactor. The validity of Eqs. [S.2] and [S.3] can be proven by equating Eqs. [S.1] and [S.2] under the assumption that 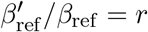 and then solving for *C*′.

### Supplementary residual *C* dependence on tissue type and 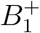

To quantitatively illustrate this effect of small residual dependence of calibration coefficient *C* on tissue type and 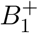 in all brains, partial Spearman correlation coefficients between the bias corrected MTsat (as an indicator of tissue type) and *C* (accounting for 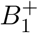 were calculated. Spearman correlations were used so as to also be sensitive to nonlinear monotonic relationships. The coefficients ranged from *ρ* = −0.532 to *ρ* = −0.749 across the brains (Table 1). This suggests that the tissue type explains up to a half of the residual 3–9% variance in the estimated calibration parameter and that the assumption is to some extent violated. We found low to medium degrees of partial Spearman correlation between 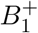 and the calibration parameters (correcting for MTsat).

## Footnotes

1 MTsat differs from the commonly used magnetization transfer ratio (MTR) (12) in that it is inherently corrected for TR and on-resonance excitation flip angle dependence (13; 37).

2 This method of computing *S*_0_ and R1 differs from that in Ref. (13), which made use of small angle approximations with respect to *α*_PD_ and *α*_T1_ to simplify the calculation. We avoid relying on the small angle approximation here because large *α*_T1_ are often used in *postmortem* high resolution imaging to impose sufficient T1-weighting (for example in our case *α*_T1_ = 84^◦^).

3 We implicitly exclude potential contributions from direct saturation of the free pool by the MT pulse (28), which would scale with *β*_loc_ but not with *m*. Both Bloch simulations of the direct saturation and fitting the experimental data with a model including a MTsat-independent term (data not shown), suggest that the direct saturation contribution is negligible (*<* 0.5 p.u.) for our off-resonant pulses.

